# Assessment of scoring functions to rank the quality of 3D subtomogram clusters from cryo-electron tomography

**DOI:** 10.1101/2020.06.23.125823

**Authors:** Jitin Singla, Kate L. White, Raymond C. Stevens, Frank Alber

## Abstract

Cryo-electron tomography provides the opportunity for unsupervised discovery of endogenous complexes in situ. This process usually requires particle picking, clustering and alignment of subtomograms to produce an average structure of the complex. When applied to heterogeneous samples, template-free clustering and alignment of subtomograms can potentially lead to the discovery of structures for unknown endogenous complexes. However, such methods require useful scoring functions to measure the quality of aligned subtomogram clusters, which can be compromised by contaminations from misclassified complexes and alignment errors. To our knowledge, a comprehensive survey to assess the effectiveness of scoring functions for ranking the quality of subtomogram clusters does not exist yet. Here, we provide such a study and assess a total of 15 scoring functions for evaluating the quality of the subtomogram clusters, which differ in the amount of structural misalignments and contaminations due to misclassified complexes. We assessed both experimental and simulated subtomograms as ground truth data sets. Our analysis shows that the robustness of scoring functions varies largely. Most scores are sensitive to the signal-to-noise ratio of subtomograms and often require Gaussian filtering as preprocessing for improved performance. Two scoring functions, Spectral SNR-based Fourier Shell Correlation and Pearson Correlation in the Fourier domain with missing wedge correction, show a robust ranking of subtomogram clusters even without any preprocessing and irrespective of SNR levels of subtomograms. Of these two scoring functions, Spectral SNR-based Fourier Shell Correlation was fastest to compute and is a better choice for handling large numbers of subtomograms. Our results provide a guidance for choosing a scoring function for template-free approaches to detect complexes from heterogeneous samples.

## 1. Introduction

Cryo-electron tomography (CryoET) has evolved as a promising tool to explore the world within a cell at molecular resolution (Guichard et al., 2010; Kürner et al., 2004; Nicastro et al., 2005). These studies have revealed the cytoskeleton organization (Chakraborty et al., 2020), assembly and disassembly of bacterial flagella motor (Kaplan et al., 2019), structures of actin networks and other cellular components (Beck and Baumeister, 2016; Gan et al., 2019; Medalia et al., 2002), membrane-associated macromolecules (Dunstone and de Marco, 2017) and native structures and organization of the cytoplasmic translation machinery, as well as nucleosome chains and filaments of the nuclear lamina in situ (Mahamid et al., 2016).

With the advancement and increased automation of CryoET, it has become easier to collect a vast amount of tomograms in a short period. Thus, we require automated methods for the analysis of these tomograms as well. Over the last few years, various efforts have been made to extract relevant information from tomograms by semi-automated and fully-automated methods. These include use of neural-networks (Che et al., 2018; Chen et al., 2017; Yu and Frangakis, 2011), template-based detection (Beck et al., 2009; Böhm et al., 2000; Lebbink et al., 2007) and template-free pattern mining (Frazier et al., 2017; Martinez-Sanchez et al., 2020; Xu et al., 2011, 2012, 2019). Template-based and neural-network-based methods are successful in detecting complexes in tomograms. However, they are limited to discover only those complexes for which structures are already known. Template-free unsupervised methods stand out as they are capable of identifying structures of unknown complexes in tomograms.

We previously developed the Multi-Pattern Pursuit (MPP) (Xu et al., 2019), which allows large-scale template-free detection of macromolecular structures in tomograms of heterogeneous samples. The method performs unsupervised clustering of subtomograms into different structural classes and uses an iterative optimization process to select the best combination of alternative clustering results. The underlying structure is then retrieved by averaging the aligned subtomograms in each cluster. MPP and all other methods based on unsupervised subtomogram clustering require an effective scoring function for robust quality assessment of clusters and filtering out of unreliable results. Such a quality score can distinguish the homogeneous and well-aligned subtomogram clusters from contaminated and misaligned clusters.

A variety of scoring functions have been developed for cryo-Electron Microscopy (cryoEM) density fitting (Vasishtan and Topf, 2011). These scoring functions measure how well the atomic structure of a complex fits into its electron density maps. Similarly, scoring functions have been used to compare the alignments between 3D electron microscopy volumes (Joseph et al., 2017). However, currently, not much attention has been devoted to scoring functions for assessing the overall quality of a subtomogram cluster, a set of aligned 3D subtomograms that likely contain the same underlying complex. Averaging these subtomograms produces the structure of the complex. The quality of subtomogram clusters depends on the alignment errors among subtomograms and whether or not all the subtomograms in a cluster contain the same underlying complex. These clusters of subtomograms could have been generated by supervised classification and alignment methods or from unsupervised (i.e., reference-free) clustering methods from cryo-electron tomograms of purified complexes, cell lysates or native cellular landscapes.

In contrast to template-based methods, clusters from unsupervised methods cannot be assessed by comparison to known template structures as the template might be unknown. So they must be evaluated by cross-comparison of the similarity of aligned subtomograms. Here, we tested 15 scoring functions and compared their ability to rank the quality of subtomogram clusters without knowledge of template structures. The quality of clusters is ranked higher when they; i) are homogenous in terms of their complex composition, and ii) constituent subtomograms are well-aligned to each other. Scoring functions were tested on sets of both simulated and experimental ground truth subtomograms. For simulated tomograms, we chose five complexes of varying size and shape from the Protein Data Bank (PDB) (Berman et al., 2000) to realistically simulate subtomograms in various different orientations and at three different SNRs (0.001, 0.01, 0.1 - Methods section). For the test on experimental subtomograms, we used a set of ∼800kDa GroEL_14_ and GroEL_14_/GroES_7_ subtomograms that have been used in other studies as quasi-standard in the field (Section 2.1).

## 2. Methods

### 2.1 Data preparation

#### 2.1.1 Simulated data

As a test set, we used five protein complexes (Table 1) with varying sizes and shapes. Atomic structures of all the five complexes were converted into density maps using the pdb2vol program in the situs package (Wriggers et al., 1999) at 0.4 nm voxel spacing and bandpass filtered at 2 nm. We generated ground truth data sets following a previously established approach for the realistic simulation of the tomographic image reconstruction process. It allows the inclusion of noise, tomographic distortions due to missing wedge, and electron-optical factors such as Contrast Transfer Function (CTF) and Modulation Transfer Function (MTF) (Beck et al., 2009; Förster et al., 2008; Nickell et al., 2005; Pei et al., 2016; Xu et al., 2019). The density maps served as input for realistically simulating the cryo-electron imaging process with a noise-factor-SNR (SNR: Signal-to-Noise Ratio) of 0.001, 0.01, 0.1 and tilt angle range ±60°. Following a well-established procedure, subtomograms were simulated with voxel size = 0.4 nm, the spherical aberration = 2.2 mm, the defocus value = -7 µm, the voltage = 300 kV, the MTF corresponding to a realistic electron detector, defined as sinc(πω/2) where ω is the fraction of the Nyquist frequency. Finally, we use a back-projection algorithm (Nickell et al., 2005) to generate a subtomogram from the individual 2D micrographs generated at the various tilt angles (Beck et al., 2009; Xu et al., 2011). For each protein complex, we generated 1000 subtomograms, each containing a randomly rotated complex. After simulation, the density values of each simulated image were normalized to zero mean and unit variance.

**Table 1:** PDB IDs of complexes used to generate clusters. First column shows PDB IDs of the target complex in the cluster and second and third column contains PDB ID of complexes with which target complex is contaminated with.

| Target complex<br>PDB ID | Contaminant<br>complex PDB IDs |  |
| --- | --- | --- |
| **1F1B** | 2BO9 | 1A1S |
| **1FNT** | 1BXR | 3DY4 |
| **2GHO** | 1QO1 | 2H12 |
| **2GLS** | 1KP8 | 1VPX |
| **2REC** | 1VPX | 1VRG |

#### 2.1.2 Experimental Data

We used experimental subtomograms previously established as benchmark sets in various studies of subtomogram alignment and classification(Förster et al., 2008; Heumann et al., 2011; Hrabe et al., 2012; Scheres et al., 2009; Xu and Alber, 2012; Yu and Frangakis, 2011). Förster et al., (2008) collected tomograms of ∼800 kDa GroEL_14_ and GroEL_14_/GroES_7_ complexes and extracted 786 subtomograms for these complexes (GroEL_14_: 214 subtomograms and GroEL_14_/GroES_7_: 572 subtomograms). We used the same set of subtomograms, which we aligned by PyTom (Hrabe et al., 2012) using the default parameters and imposed 7-fold symmetry. Out of these 572 aligned GroEL_14_/GroES_7_ subtomograms, 500 subtomograms were used to generate primary cluster for computing scores. This primary subtomogram cluster was then contaminated with GroEL_14_ subtomograms. The voxel density values were normalized with zero mean and unit variance for all the 786 subtomograms individually.

### 2.2 Generation of Subtomogram Clusters

We define a subtomogram cluster as a set of aligned subtomograms, which upon averaging, will produce the electron density map of the underlying complex. Such clusters can be produced by supervised or unsupervised clustering methods to identify and align target subtomograms. We created a large set of different subtomogram clusters of varying quality. The subtomogram cluster quality depends on the level of misalignments, i.e., the amount of alignment errors for subtomograms in a cluster and the level of contamination, i.e., the number of subtomograms in a cluster that does not contain the target complex. Contamination is a result of misclassification or clustering error. Benchmark sets of simulated subtomograms were generated for varying levels of SNRs. In the following section, we first define how misalignments and contaminations were emulated for subtomogram clusters.

#### 2.2.1 Misalignment

To generate misalignments in a subtomogram cluster, we rotated all the subtomograms in a cluster from their initial correctly aligned orientation with Euler angles that were sampled from a normal distribution ℕ(0, *sd*) with zero-mean and a defined standard deviation (*sd*). The range of rotational angles is [−180°, 180°] for each Euler angle. At a standard deviation of 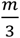 approximately 99.7% of sampled Euler angles are within the range [−*m, m*] degrees (Supplemental Figure 1). For example, a misalignment = 27 means that subtomograms were rotated in each Euler direction with angles sampled from a normal distribution 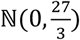, which selects ∼99.7% angles between [−27°, 27°]. In this paper, we test the scoring functions on subtomogram clusters with misalignments for each Euler angle ranging from 0 to 54 degrees.

**Figure 1:**
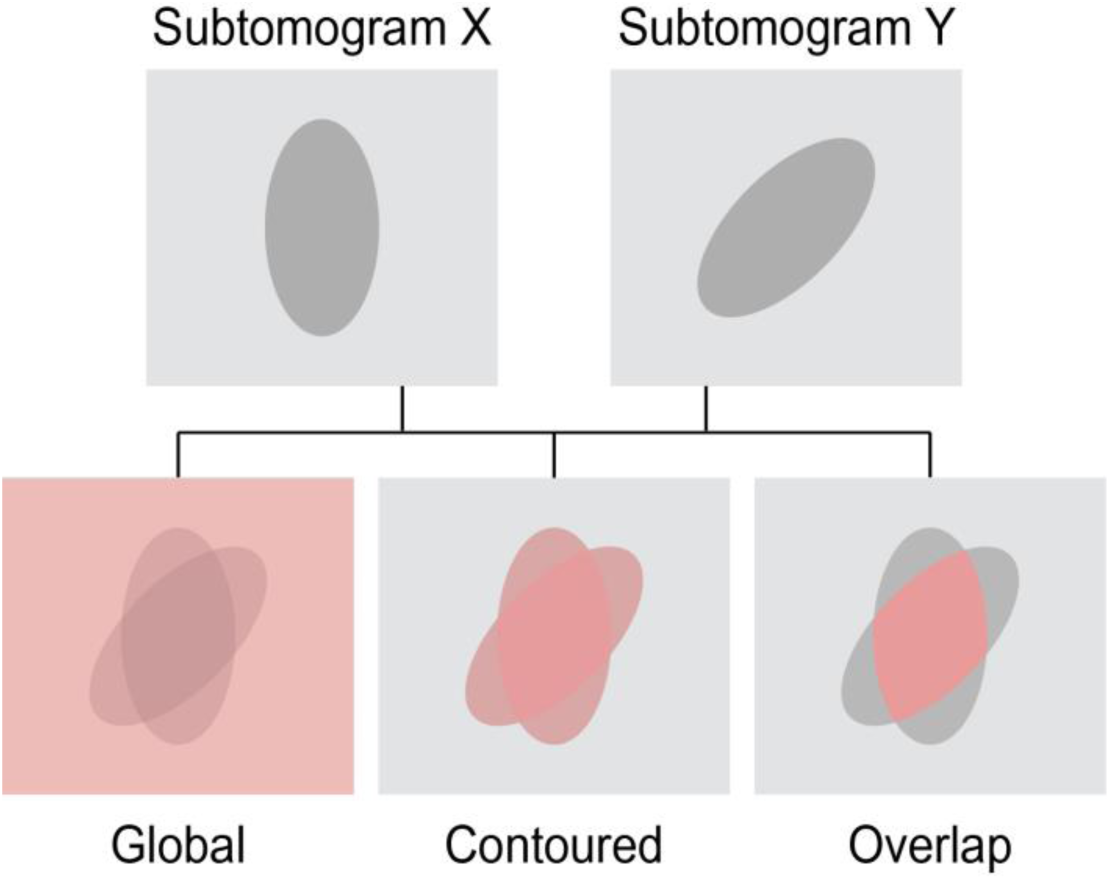
Voxel regions. Schematic representation of global, contoured and overlap regions (highlighted in red) used for computing scores between two subtomograms.

#### 2.2.2 Contamination

In both supervised classification and unsupervised clustering of subtomograms, complexes of different types but similar shapes or sizes may be falsely co-assigned to the same cluster. To assess scoring functions for their ability to detect contamination, clusters occupied with predominantly one complex *c*_*1*_ were contaminated with another complex *c*_*2*_, of similar size or shape. Clusters were generated with varying levels of contamination, defined as the percentage of the cluster size (i.e., number of subtomograms in a cluster). For instance, at contamination level *p, p*% of subtomograms in a cluster (containing subtomograms of complex *c*_*1*_) were replaced with subtomograms containing complex *c*_*2*_.

#### 2.2.3 Simulated benchmark set

For each of the five complexes, clusters were generated with misalignment values *m* ranging from [0, 54] degrees with a step of 5.4. Also, subtomogram clusters for each complex were contaminated with another complex with contamination percentage *p* ranging from [0, 40] with a step of 10. For each subtomogram cluster, we tested the assessment for contamination with two different contamination complexes. Moreover, all clusters were simulated for three different SNR = {0.001, 0.01 and 0.1} (Table 1 and Supplementary Figure 2). In total, we generated a benchmark set of 550 subtomogram clusters with varying quality in terms of misalignment and levels of contamination. Each cluster contained a total of 500 subtomograms.

**Figure 2:**
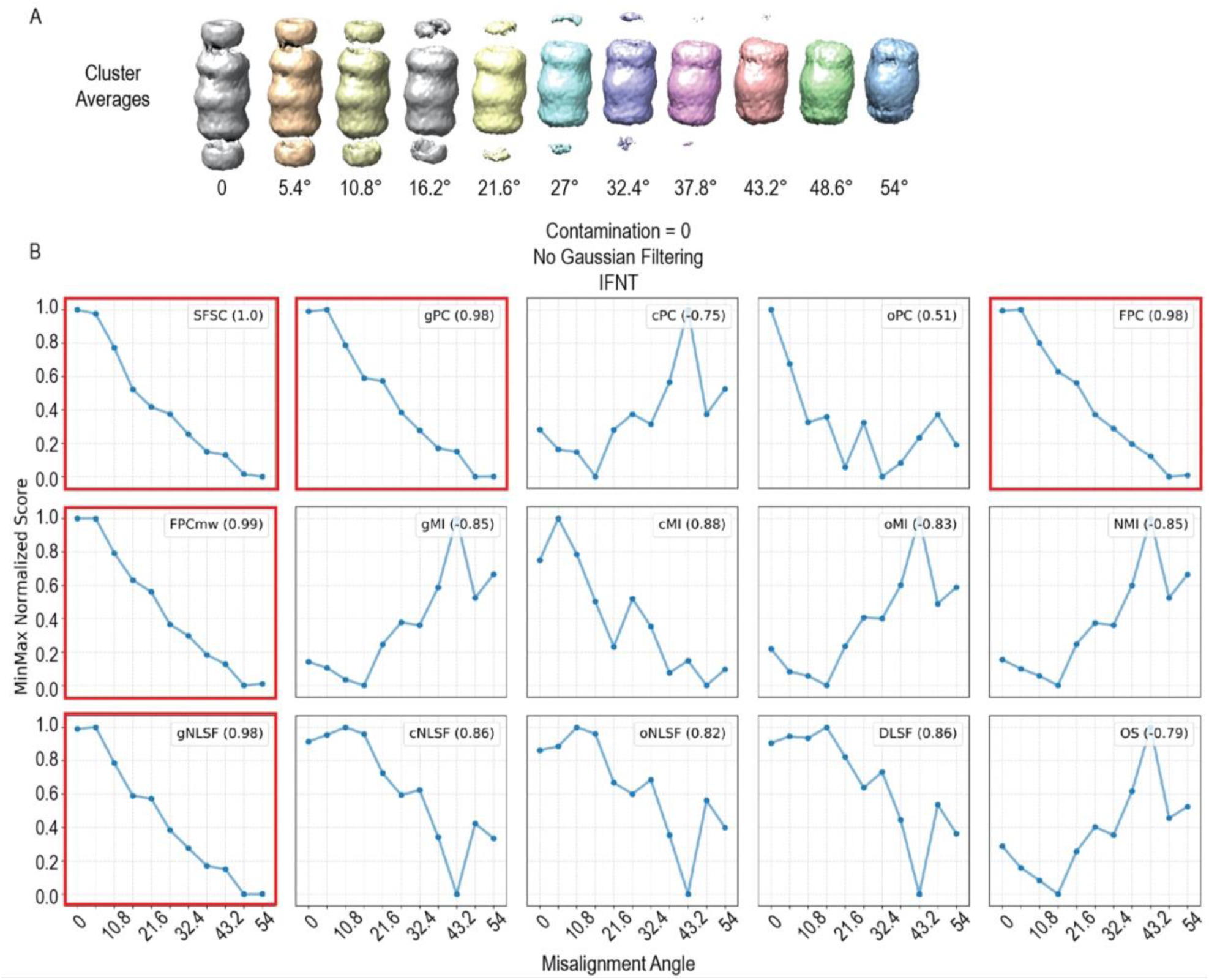
Assessment against misalignment: **A)** Cluster averages of example complex (PDB ID: 1FNT) with no contamination and with misalignment increasing from 0 (Far left) to 54 degrees (Far right). **B)** Line plots showing min-max normalized score values on y-axis varying with misalignment on x-axis for clusters constituting 1FNT alone. Legend in each subplot mentions the scoring function and its performance in Spearman’s correlation to rank clusters based on misalignment. Scores that have Spearman’s correlation above the cutoff of 0.95 are shown with subplots with red outline.

#### 2.2.4 Experimental benchmark set

Subtomogram clusters were generated for GroEL_14_/GroES_7_ using the same misalignment and contamination range as applied for simulated subtomograms. GroEL_14_/GroES_7_ clusters were contaminated with GroEL_14_. In total, a benchmark set of 55 subtomogram clusters were generated.

### 2.3 Voxel Regions

We define three different regions of voxels in a subtomogram for computing the individual scores (Figure 1).

#### 2.3.1 Global

The global score is computed from all the voxels in the subtomogram (Figure 1).

#### 2.3.2 Contoured

The contoured score is computed from a subset of voxels with density values above a threshold. We select all the voxels with density values higher than one-and-half times the standard deviation (> 1.5 *σ*). The score between two aligned subtomograms is then calculated from the union of selected voxels in both subtomograms. This step reduces the contribution of noise and focuses on voxels likely to be part of the target complex.

#### 2.3.3 Overlap

The threshold is applied as in the contoured score. The score between two aligned subtomograms is calculated from the intersection of selected voxels in both subtomograms.

### 2.4 Scoring Functions

In this section, we define the scoring functions for quality assessment of subtomogram clusters. The density values of each subtomogram image are normalized to zero mean and unit variance.

#### 2.4.1 SFSC: Spectral SNR-based Fourier Shell Correlation

SFSC measures the SNR from the variance in the voxel intensities at all spatial frequencies, as previously introduced in the MPP method (Xu et al., 2019). SFSC uses all the subtomograms in the cluster and considers missing wedge effects, one of the major distortions in cryoET, due to a limited range of angles to capture tilt series.

Say cluster *C* of size *n* contains the set of aligned subtomograms {*f*_1_, *f*_2_ … *f*_*n*_}, with Fourier Transforms {*F*_1_, *F*_2_ … *F*_*n*_} and corresponding binary missing wedge masks {*M*_1_, *M*_2_ … *M*_*n*_}. The Spectral-Signal-to-Noise Ratio (Spectral-SNR or SSNR) *η*_*r*_ at frequency *r* is defined as:

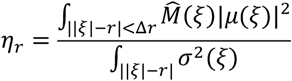

where Δ*r* = 1, *ξ* ∈ ℝ^3^ is location in Fourier space, 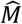 is sum of missing wedge masks:

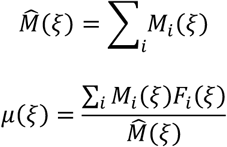

and

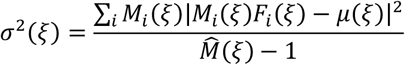

Given the SSNR (*η*_*r*_) at frequency *r*, FSC (*ζ*_*r*_) can be estimated as:

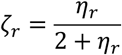

Then SFSC is defined as sum of FSC over all frequencies:

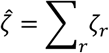

The higher the value of 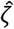 (SFSC), the higher is quality of a subtomogram cluster.

The SFSC score is computed from the set of all individual subtomograms, while all other scores are calculated from pairwise comparisons of subtomograms in the same cluster.

#### 2.4.2 gPC: Global Pearson Correlation

gPC is the *global Pearson correlation* score and uses all the voxels in both subtomograms to calculate the cross-correlation. The gPC between a pair of subtomograms (*X, Y*) is calculated as follows:

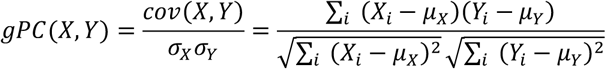

where *X*_*i*_ and *Y*_*i*_ are density values for the *i*^*th*^ voxel of subtomograms *X* and *Y*, respectively. *μ*_*X*_ and *μ*_*Y*_ are mean density values over corresponding voxel region in each subtomogram.

Because each subtomogram is normalized to zero mean and unit variance (*μ*_*X*_ = *μ*_*Y*_ = 0 and *σ*_*X*_ = *σ*_*Y*_ = 1) *ρ* = *cov*(*X, Y*). gPC is directly proportional to the cross-correlation function (CCF).

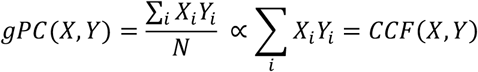

The gPC score and all following scores are calculated by randomly picking 10% of all possible pairs of subtomograms in a cluster. The total score is then defined as the average over all the pairwise scores. We show separately that for the gPC and all following scores, a random selection of 10% of pairs is sufficient to capture the population mean by comparing 10% and 50% of all possible pairs. Due to increased time complexity for computing 50% pairs (62375 pairs), we show this test for only one structure (PDB ID: 2GHO), contaminated with structures (PDB IDs: 1QO1, 2H12) at SNR = 0.01, misalignment = 21.6 degrees and contamination range [0, 30] percentage. Supplementary Table 1 shows the resulting scoring value for few scoring functions for 10% and 50% pairs. We observed that 10% of subtomogram pairs are sufficient to capture the same amount of information as 50% subtomogram pairs.

#### 2.4.3 cPC: Contoured Pearson Correlation

cPC is calculated as defined in gPC. However, only the union of voxels in both subtomograms with density values larger than the threshold (*X*_*i*_, *Y*_*i*_ > 1.5 *σ*) are considered.

#### 2.4.4 oPC: Overlap Pearson Correlation

oPC is calculated as defined in gPC. However, only the intersection of voxels from both subtomograms with density values larger than the threshold (*X*_*i*_, *Y*_*i*_ > 1.5 *σ*) are considered.

#### 2.4.5 FPC: Pearson correlation in Fourier space

We computed the Pearson correlation in the Fourier Space as well. Say *F*(*X*) and *F*(*Y*) are Fourier Transforms of subtomogram *X* and *Y* respectively. Then Pearson Correlation in Fourier space is computed as:

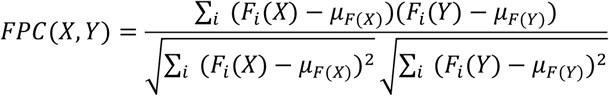

where *F*_*i*_(*X*) and *F*_*i*_(*Y*) are values at *i*^*th*^ voxel of Fourier Transforms of subtomograms *X* and *Y*, respectively. *μ*_*F*(*X*)_ and *μ*_*F*(*Y*)_ are mean intensity values of voxels in Fourier Transforms.

#### 2.4.6 FPCmw: Pearson correlation in Fourier space with missing wedge correction

We also calculated the Pearson correlation in Fourier space with missing wedge correction. The missing wedge mask in Fourier’s space is defined as the intersection of missing wedge masks of both subtomograms. Say *F*(*X*) and *F*(*Y*) are Fourier Transforms and *M*(*X*) and *M*(*Y*) are binary missing wedge masks of subtomogram *X* and *Y* respectively, then FPCmw score can be written as:

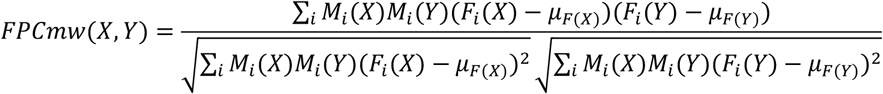

where,

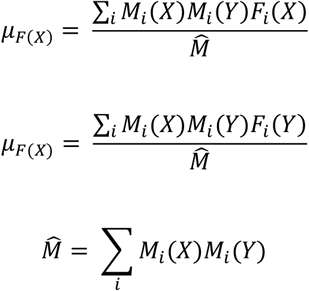

Overall we have five Pearson correlation scores computed, i.e. gPC, cPC, oPC, FPC and FPCmw.

#### 2.4.7 gMI: Global Mutual Information

Mutual information scores were previously used (i) to improve the alignment of class-averages in Single Particle Analysis (SPA) (Shatsky et al., 2009), (ii) to fit crystal structures in cryo-density maps and (iii) to assess structures determined by cryo-electron microscopy (Joseph et al., 2017; Vasishtan and Topf, 2011). Here we define a mutual information score to calculate the quality of a subtomogram cluster. The density values of all voxels in the desired voxel region were divided into *k* number of bins. The number of bins *k* was defined following the Sturges rule (Sturges et al., 1926) as:

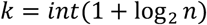

where *n* is the total number of voxels.

Marginal entropies were then calculated for both the subtomograms *X* and *Y* as

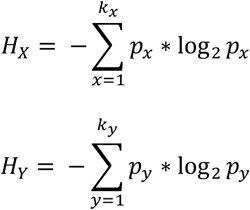

where *p*_*x*_ and *p*_*y*_ are the probabilities of finding a voxel with density values for bins *x* and *y* in the corresponding subtomograms. *k*_*x*_ and *k*_*y*_ are the number of bins into which subtomogram *X* and *Y* were divided. The joint entropy was computed as

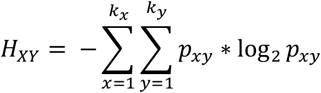

where *p*_*xy*_ is the probability of finding the pair of bins *x, y* in the aligned set of subtomograms. The joint entropy is minimum when there is no difference between subtomogram X and Y. Then gMI was calculated using all voxels in the subtomograms as:

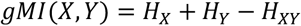

MI was calculated for all the three regions in a subtomogram as well, i.e., global, contoured, overlap. Also, if subtomograms *X* and *Y* are normalized to have zero means and unit standard deviations, *H*_*X*_ and *H*_*Y*_ are approximately equal and constant for any pair of subtomograms containing the same structure and SNR. Therefore, mutual information, in that case, is inversely proportional to joint entropy.

#### 2.4.8 cMI: Contoured mutual Information

cMI score is calculated as defined in gMI. However, only the union of voxels in both subtomograms with density values larger than the threshold (*X*_*i*_, *Y*_*i*_ > 1.5 *σ*) are considered.

#### 2.4.9 oMI: Overlap mutual Information

oMI score is calculated as defined in gMI. However, only the intersection of voxels in both subtomograms with density values larger than the threshold (*X*_*i*_, *Y*_*i*_ > 1.5 *σ*) are considered. oMI has also been used before but called Local Mutual Information (Joseph et al., 2017).

#### 2.4.10 NMI: Normal Mutual Information

We also calculated a normalized version of the mutual information sore. The NMI score is calculated as:

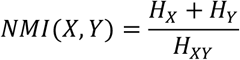

where *H*_*X*_ and *H*_*Y*_ as the marginal entropies calculated from subtomograms *X* and *Y* and *H*_*XY*_ is the joint entropy. The statistical power of estimated probabilities decreases as the overlap between subtomograms decreases. But NMI (Studholme et al., 1999) make gMI more robust to overlap volume.

#### 2.4.11 gNLSF: Global Normalized Least Square Function

Least Square Function (LSF) between two subtomograms is defined by the difference between the density values of corresponding voxels in the two aligned subtomograms.

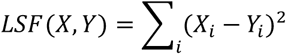

where *X*_*i*_ and *Y*_*i*_ are voxel densities at *i*^*th*^ voxel of subtomograms *X* and *Y* respectively. For global Least Square Function (gLSF), the score comes out to be directly proportional to cross correlation function.

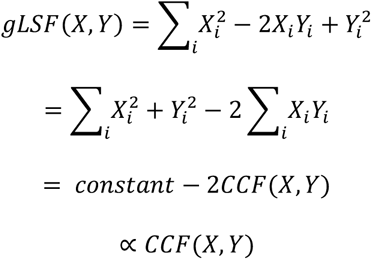

The gNLSF score is then defined by min-max normalization of gLSF and by a subtraction from 1 to define a score that increases with quality.

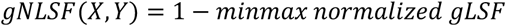

#### 2.4.12 cNLSF: Contoured Normalized Least Square Function

cNLSF score is calculated as define in gNLSF. However, only the union of voxels in both subtomograms with density values larger than the threshold (*X*_*i*_, *Y*_*i*_ > 1.5 *σ*) are considered.

#### 2.4.13 oNLSF: Overlap Normalized Least Square Function

oNLSF score is calculated as defined in gNLSF. However, only the intersection of voxels in both subtomograms with density values larger than the threshold (*X*_*i*_, *Y*_*i*_ > 1.5 *σ*) are considered.

#### 2.4.14 DLSF: Difference Least Square Function

The DLSF score is similar to LSF. However, instead of using density values, it uses the difference of density values between the pairs of corresponding voxels in the two subtomograms.

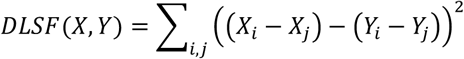

where (*i, j*) is the pair of voxels, *X*_*i*_, *X*_*j*_, *Y*_*i*_, *Y*_*j*_ are density values at voxel indices *i* and *j* for subtomograms *X* and *Y*. As the number of all possible voxel pairs can be very expensive to compute, we only used 10,000 randomly selected voxel pairs that have density values higher than a particular threshold. Here we chose that threshold to be the standard deviation of voxel densities in a subtomogram. Similar to LSF, DLSF also represents the difference between the subtomograms, so after min-max normalization of the score, we subtract it from 1. DLSF we mention throughout Results section is:

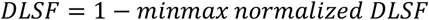

#### 2.4.15 OS: Overlap Score

The overlap score is defined as the fraction of contoured voxel regions that are part of the intersection of both subtomograms.

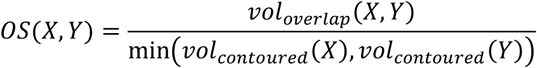

where *vol*_*contoured*_ is the volume of contoured regions in a subtomogram and *vol*_*overlap*_(*X, Y*) is the volume for overlap regions in subtomograms *X* and *Y* (contour and overlap regions are defined as previously described).

In total, we compared fifteen variations of five scoring functions (Table 2).

**Table 2:** Acronyms of all the scoring functions and their variations based on voxel regions used for computing scores (Section 2.4).

| Scoring Function | Global | Contoured | Overlap | Significant Voxels |
| --- | --- | --- | --- | --- |
| **Spectral SNR-based Fourier Shell Correlation** | SFSC |  |  |  |
| **Pearson Correlation** | gPC<br>FPC<br>FPCmw | cPC | oPC |  |
| **Mutual Information** | gMI<br>NMI | cMI | oMI |  |
| **Least Square Function** | gNLSF | cNLSF | oNLSF | DLSF |
| **Overlap Score** |  |  | OS |  |

### 2.5 Estimation of effective-SNR

#### 2.5.1 Simulated Data

We estimated the effective-SNR as previously described (Frank and Al-Ali, 1975; Xu et al., 2019). At each SNR level, we sample 10,000 pairs of aligned subtomograms for each of the five complexes. For each pair of subtomograms, we calculate the Pearson correlation of their voxel densities and then estimate a corresponding SNR according to (Frank and Al-Ali, 1975):

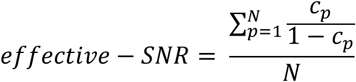

where, N is the number of pairs of aligned subtomograms and *c*_*p*_ is the Pearson correlation between subtomograms in pair *p*. To estimate the effective-SNR at given simulated SNR level, we averaged the effective-SNR for each of the five complexes. This procedure gives effective-SNR of 0.002, 0.01 and 0.08 for simulated SNR levels of 0.001, 0.01 and 0.1, respectively (Supplementary Table 2).

#### 2.5.2 Experimental Data

Similar to estimating the effective SNR for simulated subtomograms, we chose 10,000 pairs for aligned GroEL_14_/GroES_7_ experimental subtomograms and another 10,000 pairs for GroEL_14_ experimental subtomograms. The effective-SNR for GroEL_14_/GroES_7_ turns out to be ∼0.115 and for GroEL_14_ ∼0.113.

### 2.6 Gaussian Filtering of subtomograms

As a preprocessing step to score computation, subtomograms were filtered using a Gaussian filter with two kernel values (σ = 1 and σ = 2). Gaussian filtering blurs the density values in the subtomogram and emphasizes the voxels containing underlying structure while removing density variance from other voxels (Supplementary Figure 3B). We used python package Scipy to filter the 3D subtomograms (Virtanen et al., 2020).

**Figure 3:**
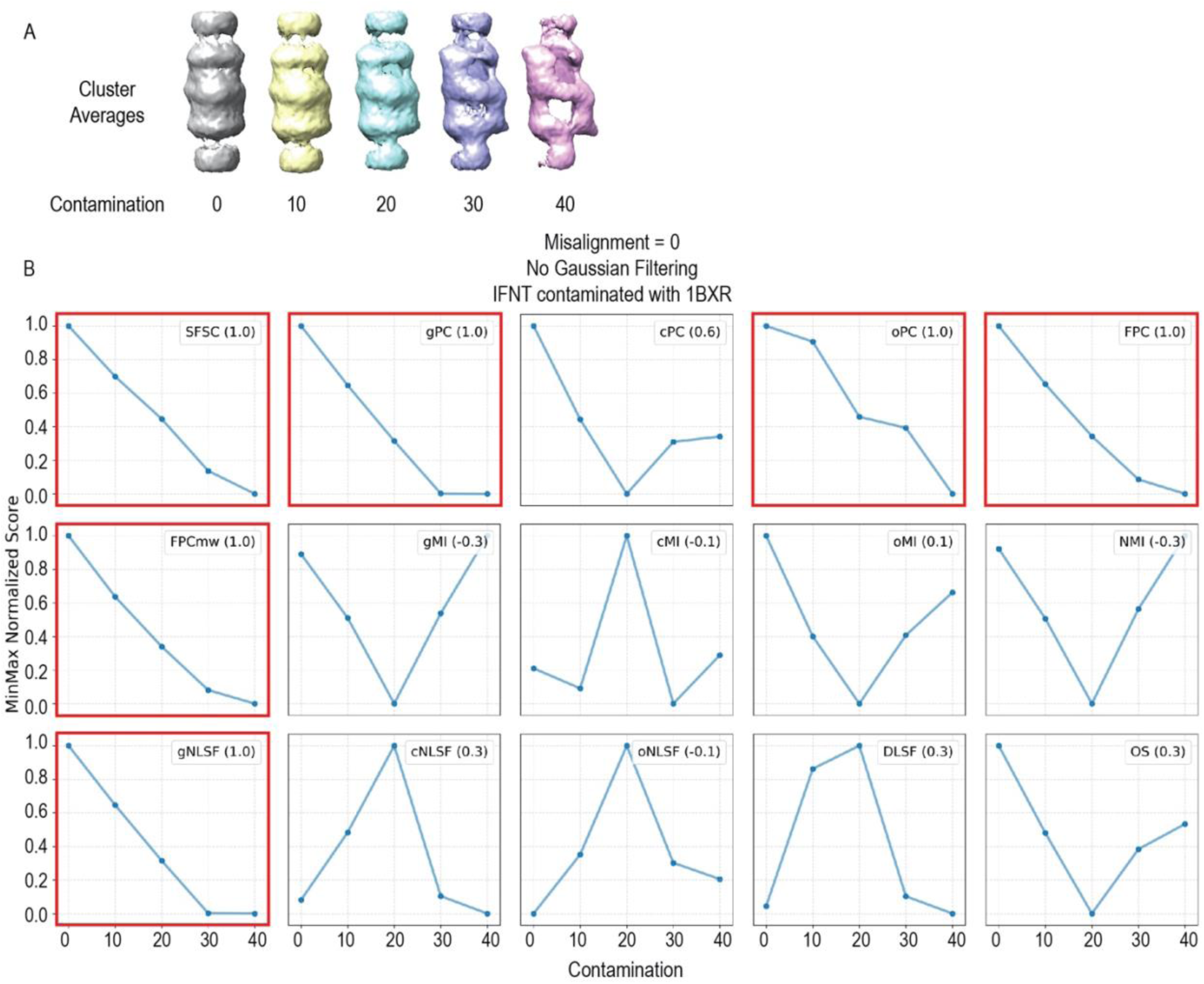
Assessment of cluster contamination: **A)** Cluster averages of example target complex (PDB ID: 1FNT) with no misalignment and contaminated with one of its assigned contaminant complex (PDB ID: 1BXR), with contamination level increasing from 0 (Far left) to 40% (Far right). **B)** Line plots shows min-max normalized score values on y-axis varying with contamination (assignment error) on x-axis for clusters constituting target complex (PDB ID: 1FNT) and contaminated with contaminant complex (PDB ID: 1BXR). Legend in each subplot mentions the scoring function and its performance in Spearman’s correlation to rank clusters based on contamination. Because we have only five sample points to compute ρ, we lower the threshold and select those functions as well-performing that have ρ > 0.85. Scores that have Spearman’s correlation above the cutoff of 0.85 are shown with subplots with red outline.

## 3. Results

The quality of a subtomogram cluster depends on various factors that include: (i) subtomogram misalignments and (ii) cluster contamination. Subtomogram misalignments (i.e., alignment errors) are non-optimal alignments of two subtomograms, which result from low accuracy in alignment programs, in particular for subtomograms of low resolution and with high noise levels. Cluster contamination (i.e., assignment error) occurs when subtomograms with structures other than the target complex are classified into the same cluster. This can be the result of errors in classification programs due to subtomograms with low resolution and higher noise levels.

To assess each scoring function for correctly ranking the quality of subtomogram clusters based on misalignment and contamination errors we compute the Spearman’s rank correlation coefficient (ρ) between the predicted subtomogram cluster quality and the amount of actual error in the clusters. Spearman’s correlation of ρ = 1 indicates that the quality score is strictly monotonic and the scoring function values decrease with increasing errors in the subtomogram clusters. The main criteria to categorize the scoring function as useful will be its ability to correctly rank the clusters in the order of their actual quality.

### 3.1 Assessment against Misalignment

We first assess the scoring function performance when only alignment errors are introduced in clusters, i.e., contamination = 0 for perfectly homogeneous clusters. Each cluster contains a total of 500 subtomograms. We generated 11 clusters for each of the five benchmark complexes, and each sampled with an increasing range of misalignments from 0 to 54° (step size = 5.4 degrees, Section 2.2, Supplementary Figure 1). Because the angles for misalignments are sampled randomly from a normal distribution, we repeated the process three times and averaged the scores over the three replicates.

Figure 2 shows each scoring function’s performance to rank the quality of clusters for an example complex (PDB ID: 1FNT) with subtomograms at SNR = 0.001 and increasing misalignment errors. The *effective-SNR* in our experimental subtomograms is estimated to be ∼0.11 (Section 2.5.2), and therefore an SNR of 0.001 for simulated subtomograms represents a challenging test case. To allow comparison, scores were min-max normalized to the range [0, 1]. To compute Spearman’s correlation (ρ), we ranked the zero misalignments as the highest rank for each scoring function.

Table 3 lists the Spearman’s correlations ρ for all scoring functions averaged over all benchmark complexes. The scoring functions differ greatly in their performance, with Spearman’s correlations ρ ranging from 1.0 to -0.93. Five scoring functions (**SFSC, gPC, gNLSF, FPC** and **FPCmw** (Section 2.4) stand out as they show excellent performance with averaged Spearman’s correlations ρ > 0.95 over the entire benchmark set, indicating that clusters can be well ranked by their ground truth quality. We noticed that all scoring functions that depend on segmented subtomogram regions (i.e., contoured and overlap regions) do not perform well for subtomograms at such low SNR value (SNR = 0.001). That is because thresholding for selecting candidate voxel regions cannot always correctly identify the volume containing the actual structure of the complex (Supplementary Figure 3A). Preprocessing can improve the thresholding for segmenting regions of the actual target complex even for very low SNR subtomograms (Section 3.3). Global and Overlap Mutual Information fail to rank clusters with subtomograms at such high noise levels. Mutual Information is inversely proportional to the joint entropy of two subtomograms containing the same underlying structure (Section 2.4.7). If subtomograms are perfectly aligned, their joint entropy is lower compared to misaligned subtomograms, i.e., the Mutual Information is higher for aligned subtomograms. This holds true only when bins with voxel intensity values of the target complex have higher probabilities than those of other regions in the subtomogram. But at very high noise levels, probabilities are more widespread across intensity bins. The performance of the mutual information score will improve by increasing the SNR of subtomograms or by preprocessing individual subtomograms. The ρ values of **gMI** and **oMI** improve when subtomograms are generated at higher SNR or after Gaussian filtering of subtomograms (Sections 3.3, 3.5).

**Table 3:** Assessment against misalignment: Spearman’s correlation (ρ) of Scoring functions vs. Misalignment for homogeneous clusters (i.e., contamination = 0). ρ for each target complex (shown with PDB ID) is mentioned separately in each column and last column shows the average ρ for scoring function over the five target complexes. Rows with bold text shows scores that performed well, i.e. had average ρ > 0.95. All values of ρ are rounded to 2 decimal places. Subtomograms were simulated at SNR = 0.001 and scores were computed without Gaussian filtering.

| Target complex | 1FNT | 1F1B | 2REC | 2GLS | 2GHO | Average $\rho$ |
| --- | --- | --- | --- | --- | --- | --- |
| **SFSC** | **1.00** | **0.98** | **0.99** | **0.99** | **0.99** | **0.99 ± 0.01** |
| **gPC** | **0.98** | **0.99** | **0.99** | **0.99** | **1.00** | **0.99 ± 0.01** |
| cPC | -0.75 | -0.54 | -0.32 | -0.51 | -0.78 | -0.58 ± 0.17 |
| oPC | 0.51 | 0.98 | 0.33 | 0.79 | 0.69 | 0.66 ± 0.22 |
| **FPC** | **0.98** | **1.00** | **0.99** | **0.99** | **0.99** | **0.99 ± 0.01** |
| **FPCmw** | **0.99** | **1.00** | **1.00** | **1.00** | **1.00** | **1.00 ± 0.00** |
| gMI | -0.85 | -0.83 | -0.72 | -0.78 | -0.93 | -0.82 ± 0.07 |
| cMI | 0.88 | 0.39 | 0.09 | 0.81 | 0.59 | 0.55 ± 0.29 |
| oMI | -0.83 | -0.75 | -0.55 | -0.70 | -0.85 | -0.73 ± 0.11 |
| NMI | -0.85 | -0.83 | -0.72 | -0.78 | -0.93 | -0.82 ± 0.07 |
| **gNLSF** | **0.98** | **0.99** | **0.99** | **0.99** | **1.00** | **0.99 ± 0.01** |
| cNLSF | 0.86 | 0.87 | 0.59 | 0.81 | 0.95 | 0.82 ± 0.12 |
| oNLSF | 0.82 | 0.74 | 0.41 | 0.76 | 0.85 | 0.71 ± 0.16 |
| DLSF | 0.86 | 0.85 | 0.59 | 0.72 | 0.93 | 0.79 ± 0.12 |
| OS | -0.79 | -0.71 | -0.43 | -0.62 | -0.79 | -0.67 ± 0.13 |

### 3.2 Assessment of Cluster Contamination

We now assess scoring functions with respect to cluster contamination, which can result from assignment errors. Clusters of a benchmark complex were contaminated with subtomograms containing other structures (Section 2.2). We generated 5 clusters per benchmark complex, which varied in the level of contamination ranging from 0 to 40% contamination. We first assess these clusters without containing any alignment errors. Figure 3 depicts the min-max normalized scores for an example complex (PDB ID: 1FNT) contaminated with another complex (PDB ID: 1BXR). Also, the scores **SFSC, gPC, gNLSF, FPC** and **FPCmw** showed the best performance in predicting the quality of the contaminated clusters (Table 4). Most scoring functions that depend on segmented subtomogram regions and scores based on mutual information fail to rank the quality of clusters accurately. This observation may be a result of the low SNR of 0.001, which reduces the quality of subtomogram thresholding and subsequently, the performance of scores relying on segmented subtomogram (Supplementary Figure 3A).

**Table 4:** Assessment of cluster contamination: Spearman’s correlation (ρ) of Scoring functions vs. Contamination for perfectly aligned clusters (i.e., misalignment = 0). ρ for each target-contaminant complex pair is mentioned separately in each column and last column shows the average ρ for scoring function over all the ten target-contaminant complex pairs. Rows with bold text shows scores that performed well, i.e. had average ρ > 0.85. All values of ρ are rounded to 2 decimal places. Subtomograms were simulated at SNR = 0.001 and scores were computed without Gaussian filtering.

| Target complex | 1FNT | | 1F1B | | 2REC | | 2GLS | | 2GHO | | Average $\rho$ |
| --- | --- | --- | --- | --- | --- | --- | --- | --- | --- | --- | --- |
| Contaminant complex | 1BXR | 3DY4 | 2BO9 | 1A1S | 1VRG | 1VPX | 1KP8 | 1VPX | 1QO1 | 2H12 |  |
| **SFSC** | **1.00** | **1.00** | **1.00** | **1.00** | **1.00** | **0.90** | **0.90** | 0.70 | **1.00** | **0.90** | **0.94 ± 0.09** |
| **gPC** | **1.00** | **1.00** | **1.00** | **1.00** | **1.00** | **1.00** | **1.00** | **1.00** | **1.00** | **1.00** | **1.00 ± 0.00** |
| cPC | 0.60 | 1.00 | 0.50 | -0.20 | 0.90 | 0.50 | 0.40 | 0.80 | 0.80 | 0.90 | 0.62 ± 0.33 |
| oPC | 1.00 | 0.90 | 1.00 | -0.70 | -0.60 | 0.80 | 0.20 | 0.50 | 0.90 | -0.70 | 0.33 ± 0.69 |
| **FPC** | **1.00** | **1.00** | **1.00** | **1.00** | **1.00** | **1.00** | **1.00** | **1.00** | **1.00** | **1.00** | **1.00 ± 0.00** |
| **FPCmw** | **1.00** | **1.00** | **1.00** | **1.00** | **1.00** | **1.00** | **1.00** | **1.00** | **1.00** | **0.90** | **0.99 ± 0.03** |
| gMI | -0.30 | 0.90 | -0.30 | -0.70 | 0.70 | -0.30 | 0.10 | 0.90 | 0.00 | 0.80 | 0.18 ± 0.57 |
| cMI | -0.10 | -0.30 | -0.70 | 0.60 | -0.90 | -0.70 | 0.70 | 0.40 | -0.50 | -0.70 | -0.22 ± 0.56 |
| oMI | 0.10 | 0.90 | 0.30 | -0.70 | 0.90 | 0.20 | 0.10 | 0.90 | 0.60 | 0.90 | 0.42 ± 0.50 |
| NMI | -0.30 | 0.90 | -0.30 | -0.70 | 0.70 | -0.30 | 0.10 | 0.80 | -0.10 | 0.80 | 0.16 ± 0.56 |
| **gNLSF** | **1.00** | **1.00** | **1.00** | **1.00** | **1.00** | **1.00** | **1.00** | **1.00** | **1.00** | **1.00** | **1.00 ± 0.00** |
| cNLSF | 0.30 | -1.00 | 0.30 | 0.90 | -0.70 | -0.10 | 0.30 | -0.90 | 0.30 | -0.30 | -0.09 ± 0.59 |
| oNLSF | -0.10 | -1.00 | -0.10 | 1.00 | -0.70 | -0.20 | 0.30 | -0.50 | 0.30 | -0.70 | -0.17 ± 0.56 |
| DLSF | 0.30 | -0.90 | 0.30 | 0.90 | -0.70 | -0.20 | -0.10 | -0.90 | 0.30 | -0.90 | -0.19 ± 0.64 |
| OS | 0.30 | 1.00 | 0.30 | -0.50 | 0.90 | 0.50 | 0.30 | 0.90 | 0.60 | 0.80 | 0.51 ± 0.42 |

### 3.3 Effect of Gaussian Filtering

Next, we test if preprocessing of subtomograms with Gaussian filtering improves the performance of scoring functions, in particular for subtomograms with low SNR values. We test Gaussian filtering with two different kernels (σ = 1 and 2, Section 2.6). Applying a Gaussian kernel enhances the global structural features of the complex against background noise for subtomograms with low SNR of 0.001 (Supplementary Figure 3B). However, with an increase in σ, naturally, the structures also lose their high-resolution features. At very low SNR (SNR = 0.001), Gaussian filtering improves the automatic thresholding of subtomograms to detect contoured and overlap regions (Section 2.3, Supplementary Figure 3A). It, therefore, improves the performances for some of the scoring functions (Table 5).

**Table 5:** Effect of Gaussian Filtering: Column 2-4: Spearman’s correlation (ρ) of Scoring functions vs. Misalignment for homogeneous clusters (i.e., contamination = 0). Column 5-7: Spearman’s correlation (ρ) of Scoring functions vs. Contamination for perfectly aligned clusters (i.e., misalignment = 0). ρ values are average over all the 10 target-contaminant complex pairs (Table 1, Supplementary Figure 2). Cells with bold text shows average ρ values that are above the cut-off, average ρ > 0.95 for misalignment and average ρ > 0.85 for contamination. All values of ρ are rounded to 2 decimal places. Subtomograms were simulated at SNR = 0.001 and Gaussian filtered with σ = 1 and 2.

| Scoring Fucntions | Against Misalignment |  |  | Against Contamination |  |  |
| --- | --- | --- | --- | --- | --- | --- |
|  | No Gaussian filtering | Gaussian kernel = 1 | Gaussian kernel = 2 | No Gaussian filtering | Gaussian kernel = 1 | Gaussian kernel = 2 |
| SFSC | **0.99** | **0.98** | 0.55 | **0.94** | **0.92** | 0.39 |
| gPC | **0.99** | **0.99** | **1.00** | **1.00** | **1.00** | **1.00** |
| cPC | -0.58 | **0.99** | **0.99** | 0.62 | **0.97** | **0.93** |
| oPC | 0.66 | **0.95** | 0.07 | 0.33 | 0.50 | 0.38 |
| FPC | **0.99** | **1.00** | **1.00** | **1.00** | **1.00** | **1.00** |
| FPCmw | **1.00** | **1.00** | **1.00** | **0.99** | **1.00** | **1.00** |
| gMI | -0.82 | -0.77 | 0.76 | 0.18 | 0.39 | 0.78 |
| cMI | 0.55 | -0.03 | 0.01 | -0.22 | -0.46 | 0.11 |
| oMI | -0.73 | **0.95** | **1.00** | 0.42 | **0.97** | **0.92** |
| NMI | -0.82 | -0.75 | 0.72 | 0.16 | 0.44 | **0.86** |
| gNLSF | **0.99** | **0.95** | **0.99** | **1.00** | 0.84 | **0.95** |
| cNLSF | 0.82 | **0.97** | **0.98** | -0.09 | 0.76 | **0.87** |
| oNLSF | 0.71 | -0.79 | -0.18 | -0.17 | -0.47 | -0.10 |
| DLSF | 0.79 | 0.91 | **0.97** | -0.19 | 0.39 | **0.86** |
| OS | -0.67 | **0.97** | **1.00** | 0.51 | **0.97** | **0.93** |

The scores (**gPC, gNLSF, FPC** and **FPCmw)**, which performed well without applying Gaussian filtering, maintain their good performance. The scores **cPC, oMI, cNLSF, DLSF** and **OS**, which failed to rank the quality of subtomogram clusters without Gaussian filter preprocessing, now show sufficiently improved Spearman’s correlation with ρ > 0.95 for assessment against misalignment and ρ > 0.85 for assessment of cluster contamination (Table 5). Therefore, these scores can rank clusters in the desired order of quality based on subtomogram misalignments and cluster contamination. In general, scores based on Mutual Information (**gMI, cMI**) fail across all Gaussian kernel settings for both misalignment and contamination tests, except for the overlap based Mutual information score (**oMI**), which shows reasonable improvements when applying a Gaussian filter (Table 5). Some scores only perform well with a narrow window of Gaussian kernel value. For instance, **oPC** (overlap Pearson correlation) performs best using Gaussian kernels with an intermediate value (σ = 1), and lose their performance with larger kernel values (Table 5). This holds true for both misalignment and contamination tests. **SFSC** decreases in performance when applying a Gaussian kernel with relatively high σ values because **SFSC** measures the variance of voxel intensities between the constituent subtomograms of the cluster (Table 5). With an increase in σ, the variation in high-frequency structural features is lost. So, **SFSC** works well when subtomograms are not preprocessed using a Gaussian filter.

### 3.4 Varying Misalignment and Contamination simultaneously

In our analysis so far, we tested scores with respect to misalignments or contamination separately. Now, we want to assess how scoring functions perform when misalignments and contaminations are introduced simultaneously. We assess the performance by calculating the average Spearman’s correlation ρ for a given score across all ten target-contaminant pairs (note that each of the five benchmark complexes is tested with two different contaminant complexes, Section 2.2, Table 1, and Supplementary Figure 2). We first calculate the Spearman’s correlation of scores for their ability to rank clusters of varying levels of misalignments at each level of contamination (from 0 to 40%) and subtomograms simulated with relatively low SNR level (SNR=0.001) (Figure 4A). We observed that **SFSC** showed excellent performance for ranking clusters against misalignment errors across all contamination levels (Figure 4A). Scoring functions based on the global Pearson correlation function (**gPC**), and its Fourier-based variants with (**FPCmw**) and without missing wedge corrections (**FPC**), and global Least Square Function (**gNLSF**) also showed excellent performance ρ > 0.95 against misalignment except at highest contamination level of 40% (Figure 4A). All other scoring functions perform very poorly in comparison.

**Figure 4:**
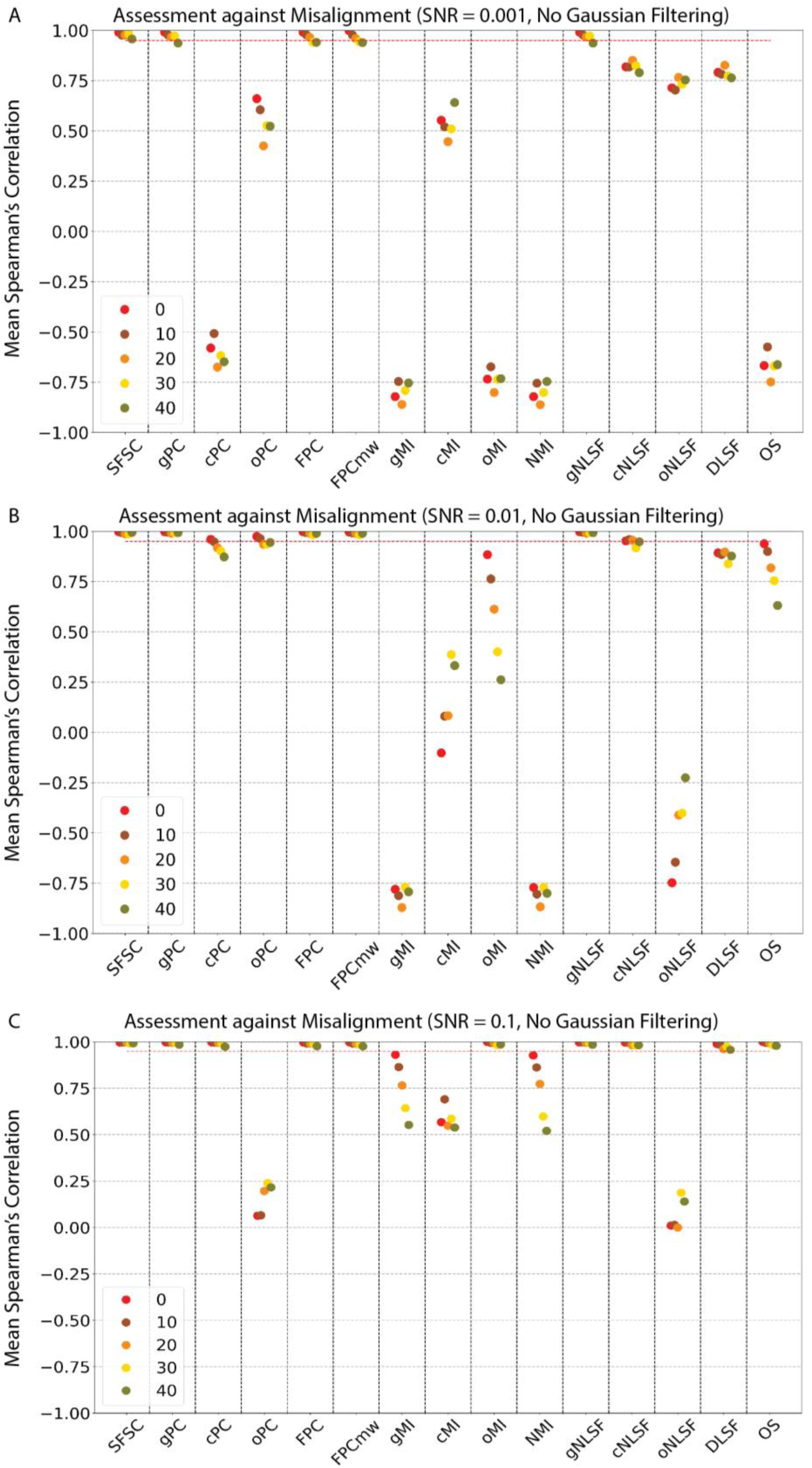
Assessment against Misalignment. Spearman’s correlation ρ (y-axis) of scoring functions (x-axis) on simulated subtomograms without Gaussian filtering. Individual panel is a scatter plot of Spearman’s correlation (ρ) of scoring functions vs. misalignment for clusters at different contamination levels. Clusters generated with the target complex can be contaminated with other complexes (Table 1, Section 2.2.2). So, each point is average ρ across all the ten target-contaminant complex pairs, except for contamination = 0, where it is averaged over only five target complexes. Red dashed line shows a cutoff value of 0.95. Subtomograms simulated at different SNR levels are shown in separate panels: **(A)** SNR = 0.001 **(B)** SNR=0.01 **(C)** SNR = 0.1.

Next, we assessed the scores for their ability to rank clusters with varying levels of contamination at each level of misalignment error ranging from 0 to 54 degrees (Figure 5A). Also here, **SFSC, gPC, FPCmw, FPC** and **gNLSF** show excellent performance to rank contamination across all levels of misalignments, except for the highest misalignment error of 54 degrees, at which **FPC** and **FPCmw** drop performance below our threshold level of ρ = 0.85. All other scores perform very poorly across all misalignment ranges and therefore, cannot rank correctly cluster quality (Figure 5A).

**Figure 5:**
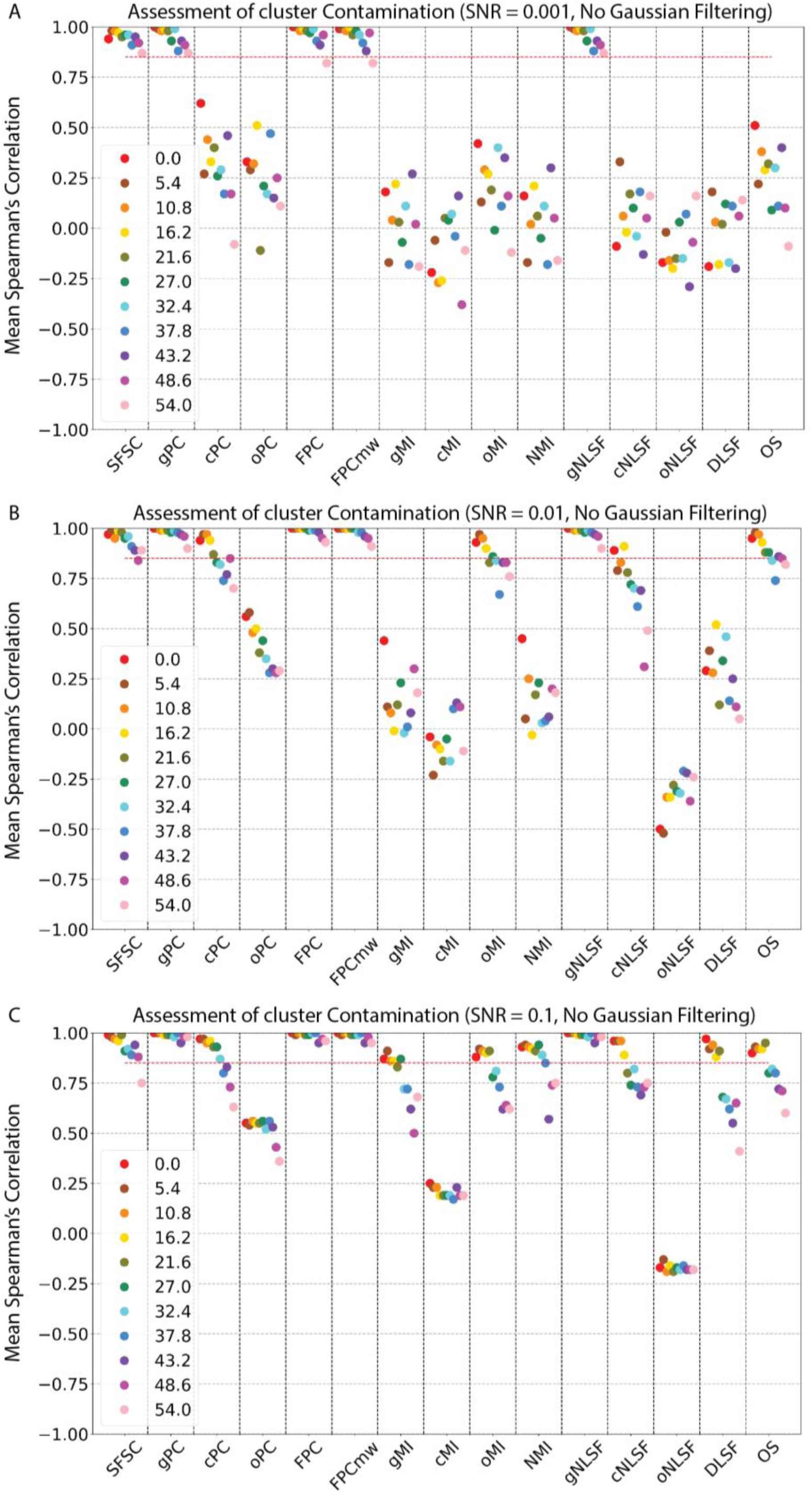
Assessment for cluster Contamination. Spearman’s correlation ρ (y-axis) of scoring functions (x-axis) on simulated subtomograms without Gaussian filtering. Individual panel is a scatter plot of Spearman’s correlation (ρ) of scoring functions vs. contamination for clusters with different misalignment errors. Clusters generated with the target complex and contaminated with other complexes can still have a varying amount of misalignment within the subtomograms (Supplementary Figure 1, Section 2.2.1). Each point is average ρ across all the ten target-contaminant complex pairs. Red dashed line shows a cutoff value of 0.85. Subtomograms simulated at different SNR levels are shown in separate panels: **(A)** SNR = 0.001 **(B)** SNR=0.01 **(C)** SNR = 0.1.

Preprocessing of subtomograms with Gaussian filters (σ= 2) improves the performance for those scoring functions that rely on segmented subtomograms. Particularly, **cPC** and **oMI** show dramatic improvements at σ = 2 for ranking misalignments across all levels of contamination even at SNR=0.001 (Supplementary Figure 4C). However, these scores perform much poorer for the ranking of contamination errors, especially when larger levels of misalignment errors are present (Supplementary Figure 4D). **cNLSF, OS** and **DLSF** scores perform better in their ability to rank clusters with varying levels of contamination only for lower levels of misalignment errors but fail to rank cluster contaminations across all levels of misalignments (Supplementary Figure 4D). Global scores based on Pearson correlations in real and Fourier space (**gPC, FPC** and **FPCmw)** retain their good performance with Gaussian filtering at high misalignment and contamination levels. Also, as seen earlier, **SFSC**’s performance slightly decreases with increasing σ in Gaussian filtering due to loss in voxel-density variations (Supplementary Figure 4). These observations confirm that **SFSC** scores work best without subtomogram preprocessing with Gaussian filters.

### 3.5 Assessment against SNR

Signal-to-Noise-Ratio (SNR) is one of the important factors that affect the performance of scoring functions. So, we simulated subtomograms at three different SNRs [0.001, 0.01 and 0.1]. Once subtomograms were simulated at a given target SNR value, we also computed the *effective-SNR* for all subtomograms as described previously (Xu et al., 2019). The *effective-SNR* levels for target SNR 0.001, 0.01 and 0.1 turn out to be 0.002, 0.01 and 0.08, respectively (Section 2.5), which indicates that the simulation process adds the required amount of noise to the subtomograms. We showed that at low SNR levels (SNR=0.001), only 5 out of 15 scoring functions are capable to rank clusters based on misalignment and contamination errors. With increasing SNR levels, we observe improved performances for scoring functions that rely on threshold-based segmentation of contoured and overlap voxel regions even without Gaussian filtering preprocessing. At the highest SNR = 0.1, almost all scoring functions (**SFSC, gPC, cPC, FPC, FPCmw, oMI, gNLSF, cNLSF, DLSF** and **OS**) show excellent performance and are all equally competent to distinguish the amount of misalignment in the clusters across all contamination levels (Figure 4C). **cPC, oMI, cNLSF, DLSF** and **OS** show excellent performance in ranking misalignments across all contamination levels (Figure 4C). However, for the ranking of contamination levels, these scores fail when high levels of misalignment errors are present (Figure 5C). **oPC** and **oNLSF** still perform very poorly across all contamination and error levels (Figure 5C).

As expected, **gMI, oMI, cMI** and **NMI** scores, which are based on mutual information, also increase in ρ with increasing SNR (Figure 4). However, at intermediate and low SNR, all three scores still fail to rank clusters reliably. At the highest SNR=0.1, only the **oMI** score reaches an acceptable Spearman’s correlation ρ > 0.95 threshold for ranking misalignments across all levels of contamination (Figure 4C). **gMI, oMI** and **NMI** can rank clusters based on contamination errors only if low levels of misalignment errors are present (Figure 5C). The **cMI** score fails to rank clusters even at the highest SNR levels. We also observed that the Overlap score (**OS**), can perform well at the highest SNR level and ranks well misalignments across all contamination levels (Figure 4C). With an improved signal component in the subtomograms, the thresholding for selecting accurate overlap voxel regions improves. So, misalignment among subtomograms can easily be recognized by the Overlap score (**OS**). Also, our complexes have non-spherical shapes, and complexes with a more spherical distribution of electron density will remain indistinguishable for overlap scores across different alignment errors. However, contaminations can only be ranked by the **OS** score when relatively low levels of misalignment errors are present (Figure 5BC).

The **SFSC, gPC, FPC** and **FPCmw** scores still outperform all other scoring functions even at high SNR levels (Figure 4 and Figure 5). Gaussian filtering with σ = 2 for subtomograms at SNR = 0.1, improves the Spearman’s correlation against contamination for many scores but only **cPC** and **oPC** show performance above our cut-off of ρ > 0.85.

### 3.6 Assessment on Experimental Subtomograms

To further assess the scoring functions with experimental subtomograms, we chose to generate clusters of GroEL_14_/GroES_7_, contaminated with GroEL_14_ (Section 2.1.2). We repeated the complete analysis with the experimental data. As it is challenging to know the exact SNR of experimental subtomograms, we estimated the *effective-SNR* (Xu et al., 2019). For aligned experimental subtomograms, the *effective-SNR* is ∼0.11 for both GroEL_14_ and GroEL_14_/GroES_7_ complexes (Section 2.5), similar to the highest SNR level of simulated subtomograms.

We observed that all scoring functions performed well for ranking misalignments across all contamination levels except **oNLSF**, which shows a weak performance similar to its performance with simulated subtomograms at SNR = 0.1 (Figure 6A). Similar to our results on simulated subtomograms, the ranking of cluster contamination across different levels of misalignments is more challenging. **gPC, FPC, gMI, oMI** and **gNLSF** rank contamination well only for clusters with relatively low misalignment errors and fail at increasing levels of misalignments. **cMI, cNLSF, oNLSF**, and **DLSF** score all show negative Spearman ranking across all levels of misalignments (Figure 6B).

**Figure 6:**
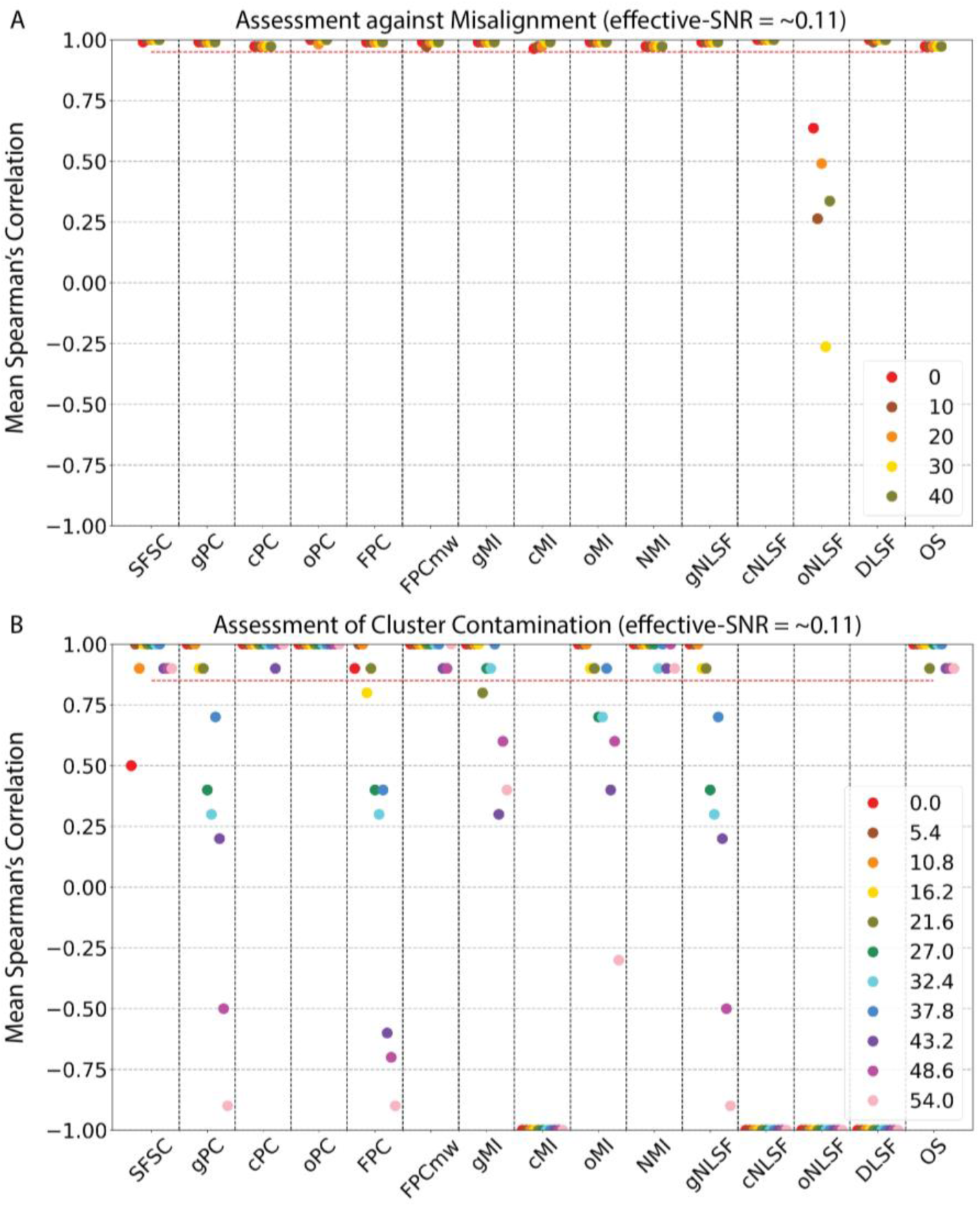
Assessment on experimental subtomograms: Scatter plot showing Spearman’s correlation (y-axis) of scoring functions (x-axis) on experimental subtomograms without Gaussian filtering. Clusters were generated using subtomograms of GroEL_14_/GroES_7_ and contaminated with GroEL_14_. Each scatter point is a ρ value. **A)** Spearman’s correlation of Scoring functions vs. Misalignment at different contamination levels. Red dashed line shows threshold value 0.95. **B)** Spearman’s correlation of Scoring functions vs. Contamination at different misalignment levels. Red dashed line shows threshold value 0.85.

Overall, **SFSC, FPCmw, cPC, oPC, NMI** and **OS** show excellent performance above the threshold of ρ > 0.85 also for ranking contamination at various levels of misalignments (Figure 6B). It is interesting to note that the Fourier-based Pearson correlation score works well only with missing wedge corrections (**FPCmw**) but performs much worse without missing wedge corrections (**FPC**) at high levels of misalignments.

### 3.7 Time Complexity

The time complexity of scoring functions varies based on the type of computations required. Fourier Space-based scores require computing the Fourier Transform of each subtomogram, whereas all the mutual information variants need to bin the voxel densities first. The time complexity reported here is on a single-core machine, with all the 500 (cluster size) subtomograms and 500 masks loaded in the memory, that is, I/O operations are not included in the time complexity measurements. Gaussian filtering or any other preprocessing step increases the time complexity further. Table 6 shows the time required to compute each score without Gaussian filtering for a cluster with 500 subtomograms. **SFSC** shows the best computational efficiency and is computationally more efficient by orders of magnitudes compared to almost all other scores. **gPC** and **gNLSF** are linearly proportional to one another without Gaussian filtering and produce the same results, but **gNLSF** takes one-fifth the time required by **gPC**, so **gNLSF** can be used instead of **gPC** to save time. As all the pairwise computations are independent, calculations are parallelizable on multi-core machines. **SFSC** can also be computed in parallel (Xu et al., 2019).

**Table 6:** Time complexity: Time required to compute score values on cluster size of 500 subtomograms and with subtomogram and mask of size 91_3_ voxels. Time was computed without Gaussian filtering, on single core computer and with all the files already loaded in the memory.

| Score | Time (in s) |
| --- | --- |
| **SFSC** | **28.70** |
| gPC | 261.84 |
| cPC | 497.96 |
| oPC | 426.82 |
| FPC | 1047.05 |
| FPCmw | 985.62 |
| gMI | 1353.70 |
| cMI | 1584.96 |
| oMI | 1689.41 |
| NMI | 1369.82 |
| gNLSF | 52.91 |
| cNLSF | 476.84 |
| oNLSF | 408.29 |
| DLSF | 13190.03 |
| OS | 380.24 |

## 4. Discussions

We compared fifteen scoring functions to test their ability to rank the quality of subtomogram clusters, which vary in the amount of misalignment and contaminations errors. Such clusters can readily be generated by unsupervised clustering methods from tomograms containing a heterogeneous set of complexes. Misalignment errors are a result of non-optimal alignments of subtomograms to each other and contaminations are a result of assignment errors, which assign subtomograms containing non-target complexes to subtomogram clusters. We require that a well-performing scoring function should be able to rank or distinguish clusters in their order of quality according to the amount of misalignment and contamination errors. We assessed the scoring functions over a wide range of SNR levels for simulated subtomograms and on an experimental benchmark dataset.

Overall, we observe a large variation in the performance of scoring functions. Only five scoring functions (**SFSC, gPC, FPCmw, FPC** and **gNLSF**) can rank clusters robustly across all variable conditions and SNR levels.

We found that **Spectral SNR-based Fourier Shell Correlation (SFSC)** showed the best performance to rank alignment as well as contamination errors across all conditions without the need for subtomogram preprocessing. Moreover, **SFSC** shows other advantages. Its computation was the fastest among all the scoring functions, in some cases by several orders of magnitudes. **SFSC** is directly computed from all the subtomograms of the cluster and therefore does not require computation of pairwise scores from randomly selected pairs of subtomograms. That means it is free from potential biases from a limited sampling of all pairwise combinations of subtomograms when calculating the quality score of large clusters. Moreover, SFSC performs well for subtomograms with low SNR levels even without Gaussian blurring of subtomogram.

Scores based on global Pearson correlation (**gPC, FPCmw** and **FPC**) show robust performance for both against misalignment (ρ > 0.95) and contamination (ρ > 0.85) across all conditions, while Pearson’s correlation scores based on segmented/contoured subtomograms (**cPC, oPC**) fail for subtomograms at low SNR levels, in particular for ranking clusters based on contaminations while also containing larger levels of misalignment errors. Preprocessing with Gaussian filtering can improve their performance, but not to a sufficient level for robustly ranking these clusters. Although Pearson correlation scores with (**FPCmw**) and without missing wedge correction (**FPC**) can both rank experimental clusters based on misalignments, only missing wedge corrected scores (**FPCmw**) can rank clusters on contamination over all levels of misalignments (Figure 6B). **SFSC** and **FPCmw** utilize missing wedge information (Section 2.4.1 and 2.4.6), which gives these scores an advantage by constricting their computation to only valid missing wedge masks and ignoring frequency regions that are missing in missing wedge mask.

### Scores based on contoured subtomograms are unreliable at lower noise levels

Scores based on mutual information are highly sensitive to SNR levels and fail to rank simulated clusters at low SNR levels. Among all mutual information-based scores, only the **overlap Mutual Information (oMI)** performs above our threshold (ρ > 0.95) for ranking misalignments, but only at the highest SNR level of SNR = 0.1 or after Gaussian filtering with σ = 2 for subtomograms with lower SNR. In previous studies (Joseph et al., 2017; Vasishtan and Topf, 2011), mutual information-based scores performed much better when applied to the ranking of atomic structures fitted into density maps from cryo-electron microscopy. This is because the 3D volumes used in previous studies have density values concentrated on the target complex regions, i.e., almost no noise component in the 3D-EM volumes. Because mutual information uses the probability of density values in different bins, applications without high noise levels generate distinct probability profiles. In our analysis of subtomograms containing high noise levels and missing wedge effects, mutual information fails unless we focus only on the overlap regions, which ensures density values are considered only from voxels that contain the target complex. This is only reliably possible either at high SNR level or after Gaussian filtering. Like most other scores, **NMI** was able to rank the experimental subtomograms with a relatively high SNR level.

We also conclude that preprocessing of subtomograms with Gaussian filters improves the performance of some scoring functions that depend on contoured and overlap voxel regions and decreases the performance of scores like **SFSC** that are dependent on the global variation of voxel intensities. Applying Gaussian filters to all subtomograms adds further to the time complexity. Moreover, scoring functions like **oPC** perform well only in a certain window of Gaussian filtering, which introduces uncertainty in determining the optimal σ value when performing a quality assessment of subtomogram clusters. So using scores that perform well without Gaussian filtering seems to be a better choice.

## Conclusion

With increasing subtomogram SNR, the performance of most scoring functions improves. Because it is always challenging to determine the exact SNR level of subtomograms, it is recommended to choose a scoring function that performs robustly over a wide range of SNR levels and ideally without the need for preprocessing. Although it will require further testing and analysis, our analysis shows that SFSC might be a useful choice for determining the cluster quality of 3D images that are prone to high noise levels. Also it is the fastest method and performs well without the need for preprocessing. In general, the choice of the scoring function is dependent on the context and goal of the problem statement. We believe the analysis done in this paper, will help users to choose a relevant function for their problems, as we move towards unsupervised methods in cryo-electron tomography.

## Acknowledgments

We thank High Performance Computing cluster resources available at Quantitative and Computational Biology department at University of Southern California, so we could run massive number of computations in parallel over 500 CPU cores. This work was supported by NIH R01GM096089, Arnold and Mabel Beckman Foundation (BYI), NSF career 1150287 to F.A.

## Author contributions

J.S. and F.A. conceived the study. J.S. implemented the code and ran analysis with inputs from F.A., K.W. and R.C.S. J.S. and F.A. analysed the results and wrote the paper with comments from K.W. and R.C.S.

## Supplementary Figures and Tables

**Supplementary Figure 1:**
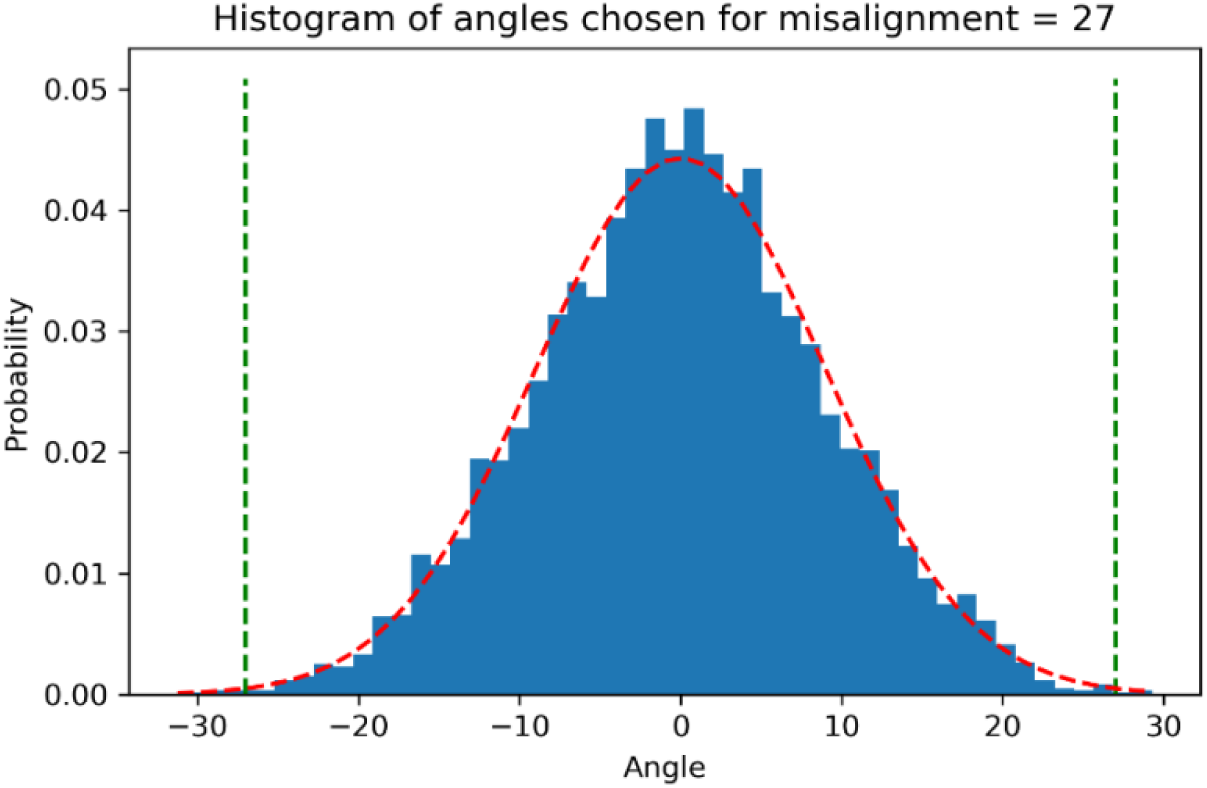
Example histogram of 5000 angles selected for misalignment = 27 degrees. Angles are selected from normal distribution 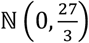 i.e., a zero-mean normal distribution with s.d. of 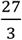. Choosing s.d. of 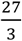, makes sure we choose approximately 99.7 percent of angles within the range [−27°, 27°]. Green dash lines mark −27° and 27° covering 99.72% of angles and red dashed curve shows the ground Gaussian distribution.

**Supplementary Figure 2:**
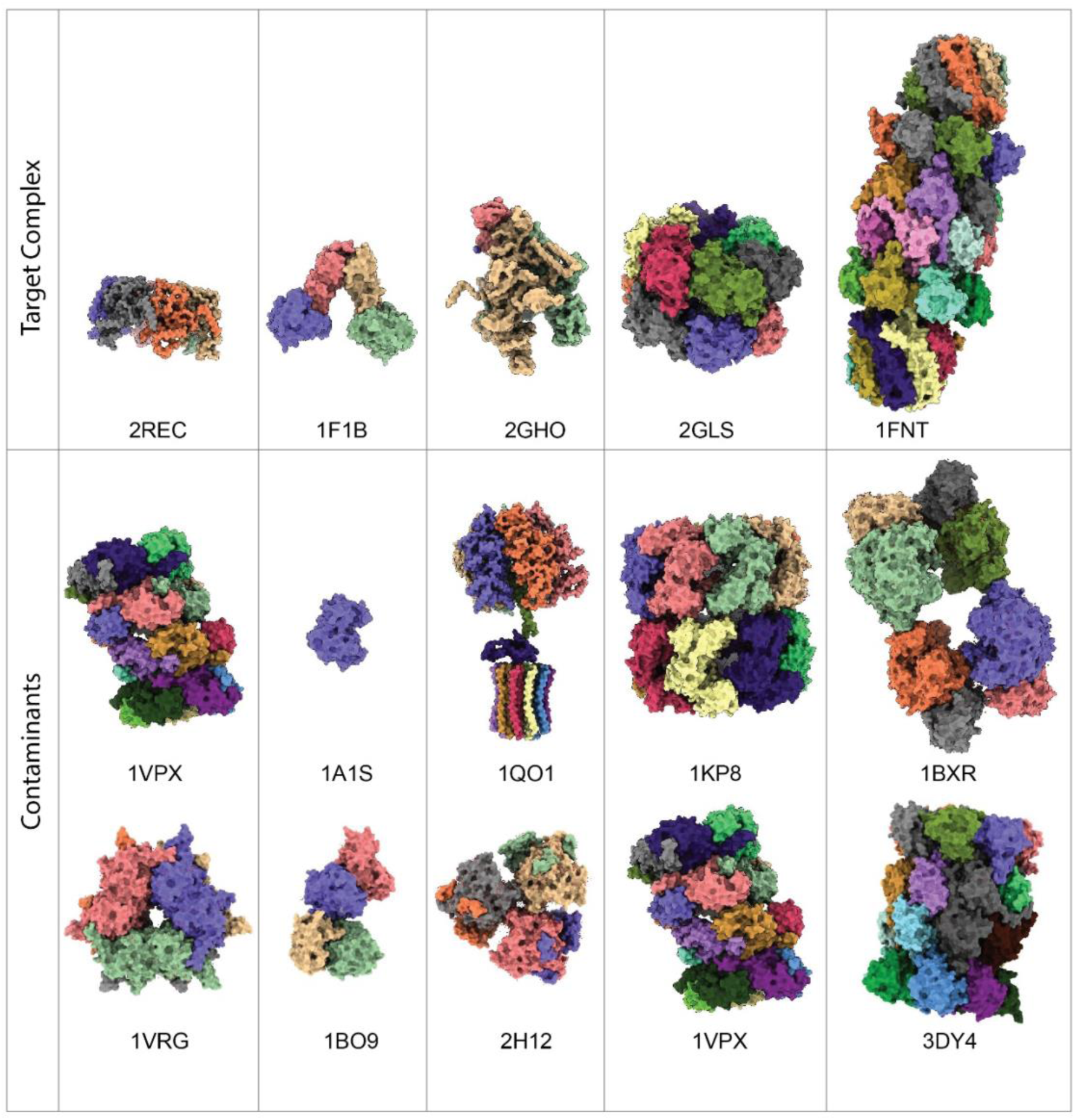
Structures of target complexes used for generating clusters and contaminant complexes used for contaminating the clusters of target complexes. PDB IDs of each complex are mentioned below the structure (Table 1).

**Supplementary Figure 3:**
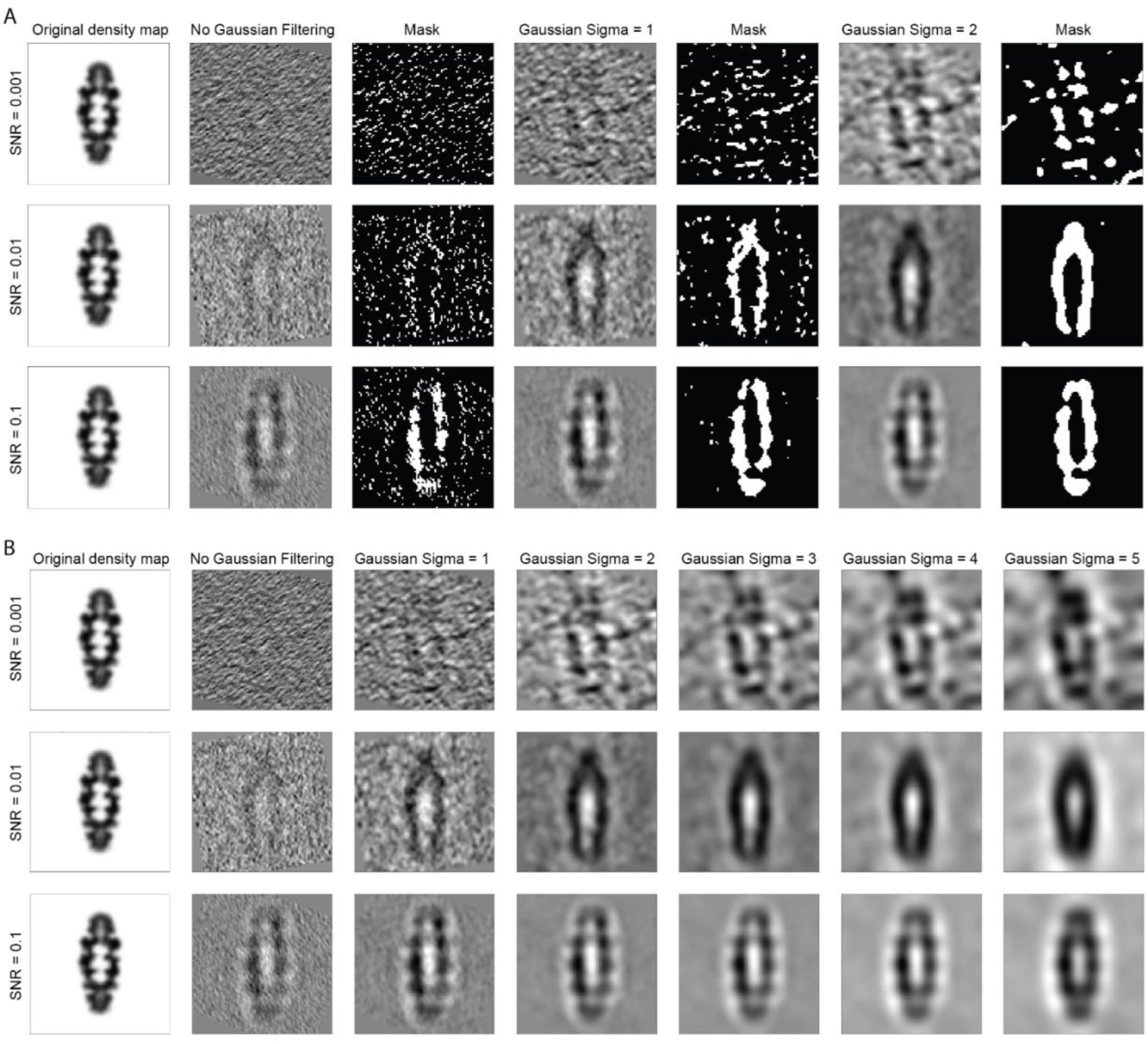
**A)** Segmented masks of subtomograms simulated for example complex (PDB ID: 1FNT) with varying SNR levels and Gaussian filtered with σ = 1 and 2. Only center slices of subtomograms and segmented masks are shown. **B)** Center slice of subtomograms simulated for example complex (PDB ID: 1FNT) with varying SNR levels and Gaussian filtered with σ in range 1 to 5.

**Supplementary Figure 4:**
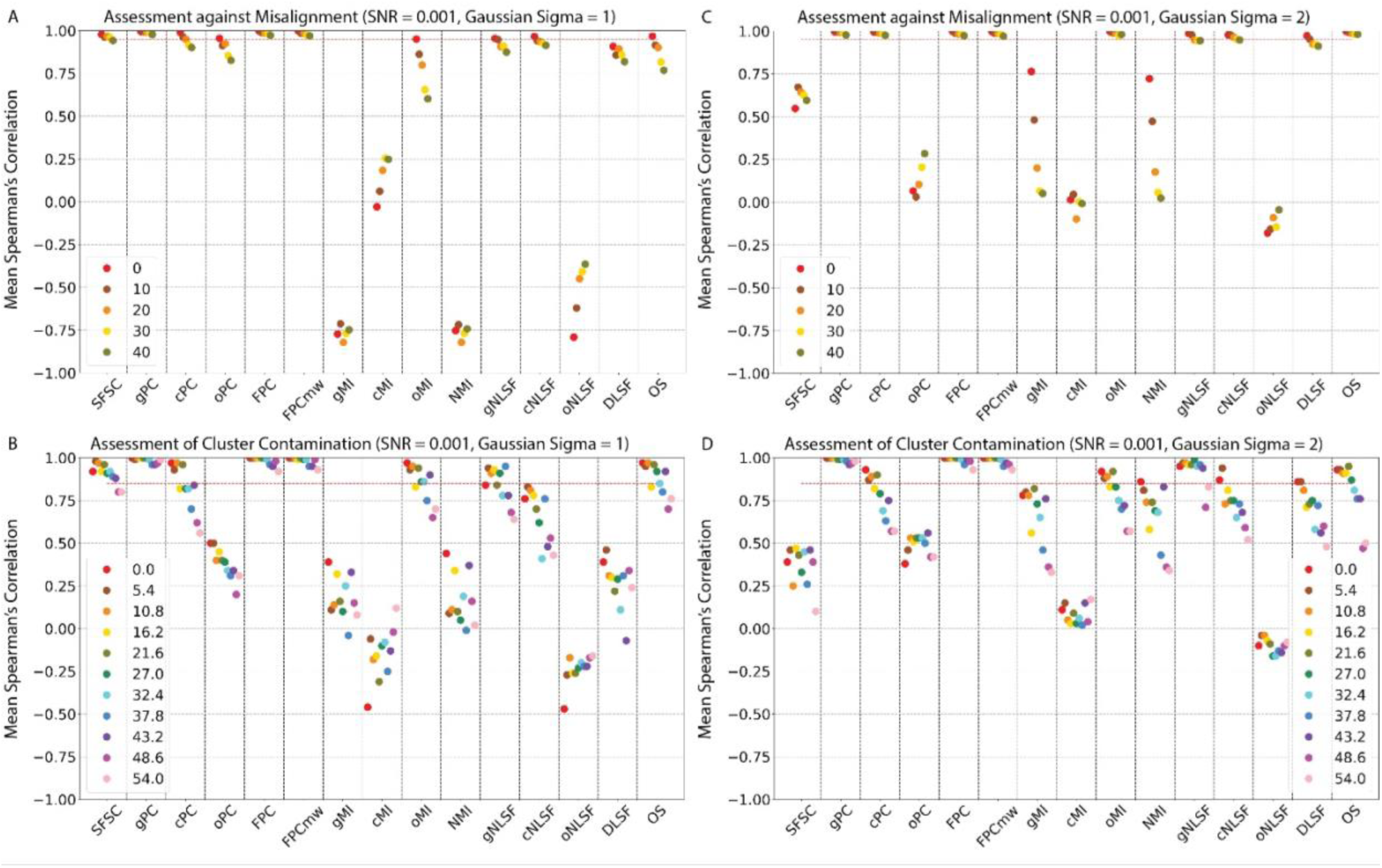
Assessment against misalignment and contamination simultaneously with Gaussian Filtering at SNR = 0.001: Scatter plot showing Spearman’s correlation ρ (y-axis) of scoring functions (x-axis) on simulated subtomograms at SNR = 0.001 and Gaussian filtered with σ = 1 and 2. Each point is average ρ across all the ten target-contaminant complex pairs. **(A, C)** Spearman’s correlation ρ of scoring functions vs. misalignment at different contamination levels. Red dashed line shows threshold value 0.95 (A: σ = 1, C: σ = 2). **(B, D)** Spearman’s correlation of scoring functions vs. contamination at different misalignment levels. Red dashed line shows threshold value 0.85 (B: σ = 1, D: σ = 2).

**Supplementary Table 1:** Absolute score values for few of the scoring functions before min-max normalization computed for pair 2GHO (target complex) and 1QO1 (contaminant complex) at SNR 0.01, misalignment = 21.6 degrees and contamination range [0, 30]. Within each contamination column, sub column shows scores computed for either 10% (12,475 pairs) or 50% (62,375 pairs) of randomly selected subtomogram pairs from all possible subtomogram pairs (124,750 pairs) from cluster of size 500 subtomograms.

|  | Contamination | 0 |  | 10 |  | 20 |  | 30 |  |
| --- | --- | --- | --- | --- | --- | --- | --- | --- | --- |
|  | Percentage of Pairs | 10 | 50 | 10 | 50 | 10 | 50 | 10 | 50 |
| Scores | gPC | 0.008 | 0.008 | 0.007 | 0.007 | 0.006 | 0.006 | 0.005 | 0.005 |
|  | cPC | -0.629 | -0.629 | -0.631 | -0.631 | -0.633 | -0.633 | -0.634 | -0.634 |
|  | oPC | 0.059 | 0.059 | 0.052 | 0.051 | 0.044 | 0.045 | 0.039 | 0.04 |
|  | FPC | 0.011 | 0.011 | 0.009 | 0.009 | 0.007 | 0.008 | 0.006 | 0.006 |
|  | FPCmw | 0.015 | 0.015 | 0.012 | 0.013 | 0.01 | 0.011 | 0.009 | 0.009 |
|  | gMI | 0.048 | 0.048 | 0.047 | 0.048 | 0.047 | 0.047 | 0.046 | 0.048 |
|  | oMI | 0.058 | 0.058 | 0.057 | 0.057 | 0.057 | 0.057 | 0.056 | 0.057 |
|  | OS | 0.093 | 0.093 | 0.092 | 0.092 | 0.091 | 0.092 | 0.091 | 0.091 |

**Supplementary Table 2:** *effective-SNR* for each of the five benchmark complexes computed for different simulated SNR levels. Last two rows show average and rounded *effective-SNR* for each SNR level.

| PDB | Simulated SNR |  |  |
| --- | --- | --- | --- |
|  | 0.1 | 0.01 | 0.001 |
| 1F1B | 0.0792 | 0.0105 | 0.0023 |
| 1FNT | 0.0655 | 0.0093 | 0.0014 |
| 2GHO | 0.0747 | 0.01 | 0.0025 |
| 2GLS | 0.0731 | 0.0093 | 0.002 |
| 2REC | 0.0819 | 0.0104 | 0.0017 |
| **effective-SNR** | **0.0791** | **0.0099** | **0.0018** |
| **After Round-off** | **0.08** | **0.01** | **0.002** |

